# An alternative explanation for reported integration and competition between space and time in the hippocampus

**DOI:** 10.1101/2025.03.10.642114

**Authors:** Federico Szmidt, Camilo J. Mininni

## Abstract

Hippocampal place and time cells are thought to be part of the brain’s spatial and temporal representation. In the article “Integration and competition between space and time in the hippocampus”, Chen *et al*. showed that hippocampal neurons with mixed selectivity for space and time shift their firing fields depending on the animal’s speed. Here, we reproduce this phenomenon with a simple continuous line attractor that only integrates the animal’s velocity. Since our model has no genuine time-encoding capabilities, it constitutes an alternative explanation for Chen *et al*. findings and challenges the claim that the observed firing field shifts are sufficient evidence for a competitive and integrated representation of space-time, as the authors suggested.

---

The hippocampus is a key brain structure for space and time encoding. In the CA1 subfield, place cells activate when the animal occupies a specific location in its environment[1]. Analogously, time cells activate at a specific time within a given time frame[2]. Together, these two neural populations are believed to be part of the neural system that allows animals to be situated in space and time[3]. However, little is known about how these two neural populations might interact. Recentrly, Chen and colleagues[4] studied the interaction of place and time encoding in the mouse hippocampus. They recorded the activity of CA1 neurons while mice had to run in 1-dimensional tracks at different velocities, and found that a high proportion of the recorded neurons were simultaneously place and time cells. Notably, the firing field of these mixed-selectivity neurons shifted depending on the animal’s speed: at higher speeds the time fields retreated to earlier times, and the place fields shifted towards more distant positions. Moreover, the authors described a trade-off between space and time information encoding, in which the more a cell encoded time, the less it encoded position, and vice-versa. The authors concluded that there is an integrated and competitive representation of time and space taking place in CA1. While we believe that the experiments performed by Chen et al. are valuable, we think there are considerable reasons to question the conclusions they have arrived at. For starters, the choice of a task where animals move in only one direction is problematic, because it makes space and time inevitably correlated. To better understand this, we can think of a pure place cell with no information regarding time. For this cell, we would expect to see no time field at all. This is what we would find in an animal randomly exploring an open field. However, if the animal always moves in the same direction, as in Chen et al., then a time field is expected, since position and time are correlated. If an animal moves at constant speed, then a pure place cell will have an apparent time field at time *p*_*n*_*/v*, where *p*_*n*_ is the animal’s position when the neuron activates. This time field recedes as velocity increases. Conversely, a pure time cell will fire at position *t*_*n*_*v*, where *t*_*n*_ is the time at which the time cell activates, and the apparent place field will move forward as velocity increases. Apparently, the shift’s direction in Chen et al. can be explained purely based on the task design, suggesting that the shifts are not a true feature of the neurophysiology of space-time encoding.

To make the above reasoning more concrete, we implemented a simple continuous line attractor[5] that only integrates the animal’s velocity and, therefore, lacks genuine time-encoding neurons to draw conclusions regarding the nature of joint space-time encoding. The model is characterized by a variable *n*(*t*) and its derivative with respect to time *dn*(*t*)*/dt* (the line attractor’s velocity). We assumed that each position in the line attractor (the state *n*) represents a distinct active neuron, that the attractor’s velocity is a function of the animal’s velocity *v*(*t*), that *n* is set to zero at the start of a lap, and that *v*(*t*) is always positive, so as *dn/dt*. Therefore, the modelled animal only moves in one direction (as in Chen et al.), and the line attractor tracks its displacement. If *dn/dt* = *v*(*t*) the system integrates velocity perfectly, and each position in space can be mapped back to a neuron in the line attractor. Hence, neurons are perfect place cells. However, if *dn/dt* = *f* (*v*), being *f* a concave, positive, non-decreasing function (Fig. 1A), then it can be proved (see *Methods*) that firing fields shift with velocity just like in Chen et al. If we choose *dn/dt* to be *f* (*v* · *η*), we can use parameter *η* to control the concavity of *dn/dt*. If *η* is small, *f* approximates the identity function for a suitable range of velocities, and neurons tend to be place cells (Fig. 1 B-C). With intermediate values of *η*, we have the case of Fig. 1 D-E, in which none of the neurons is a pure time or place cell, and both types of firing field shift (compare with Figure 4B. in Chen et al.). If *η* is high, neurons tend to become pure time cells (Fig. 1 F-G). Therefore, the shorter the shifts are in one dimension, the longer they are in the other. This replicates the trade-off reported in Figure 6 of Chen et al. (see also Fig. 4). The authors acknowledge the possibility that their results could be explained by a model that only considers place fields combined with speed (a pure place cell model (PPC)). To tackle this issue, they computed a Spatial Selectivity Index (SSI) and a Time Selectivity Index (TSI), finding that observed TSI values were systematically higher than the TSIs predicted by the PPC model. Its is worth noting that the range of selectivity indices was wide, and both indices were positively correlated, in apparent contradiction with the trade-off of space-time information. To explain this correlation, we added noise to our model and assumed that neurons in the population differ in the variance of their activity. Neurons with high variances will have wide firing fields (for *both*, time and space dimensions), leading to lower SSI and TSI values. This explains the positive correlation between indices (Fig. 2; compare with Chen et al. Figure 2 H-J), while the difference between TSI and the TSI under the PPC model is a consequence of the space-time information trade-off. In the same vein, authors found that a generalized linear model (GLM) that takes space and time as predictors to explain neural activity was better (lower Bayesian Information Criterion value (BIC)) than a model that takes space and speed. They found this result as another indication of a genuine space-time interaction. We employed our model with noise to generate neural activation events and fitted GLMs as in Chen et al. Our model displayed lower BIC levels for the space x time model, replicating the median values reported in Chen et al. (line attractor model: ΔBIC_*time*−*speed*_ = −4.2 · 10^−4^; in Chen et al.: ΔBIC_*time*−*speed*_ between −7.0 · 10^−3^ and −1.5 · 10^−3^ median value). Finally, in the analysis shown in Figure 5 of Chen et al., the authors found a non-linear dependence of space and time with speed: at higher speeds place fields shifted less than expected under a linear hypothesis. Again, our simple model reproduces the experimental results (space difference: observed-predicted = −2.5 cm (−1.3 cm median value in Chen et al.); time difference: observed-predicted = 0.78 s (0.25 s median value in Chen et al.)).

**Figure 1.**
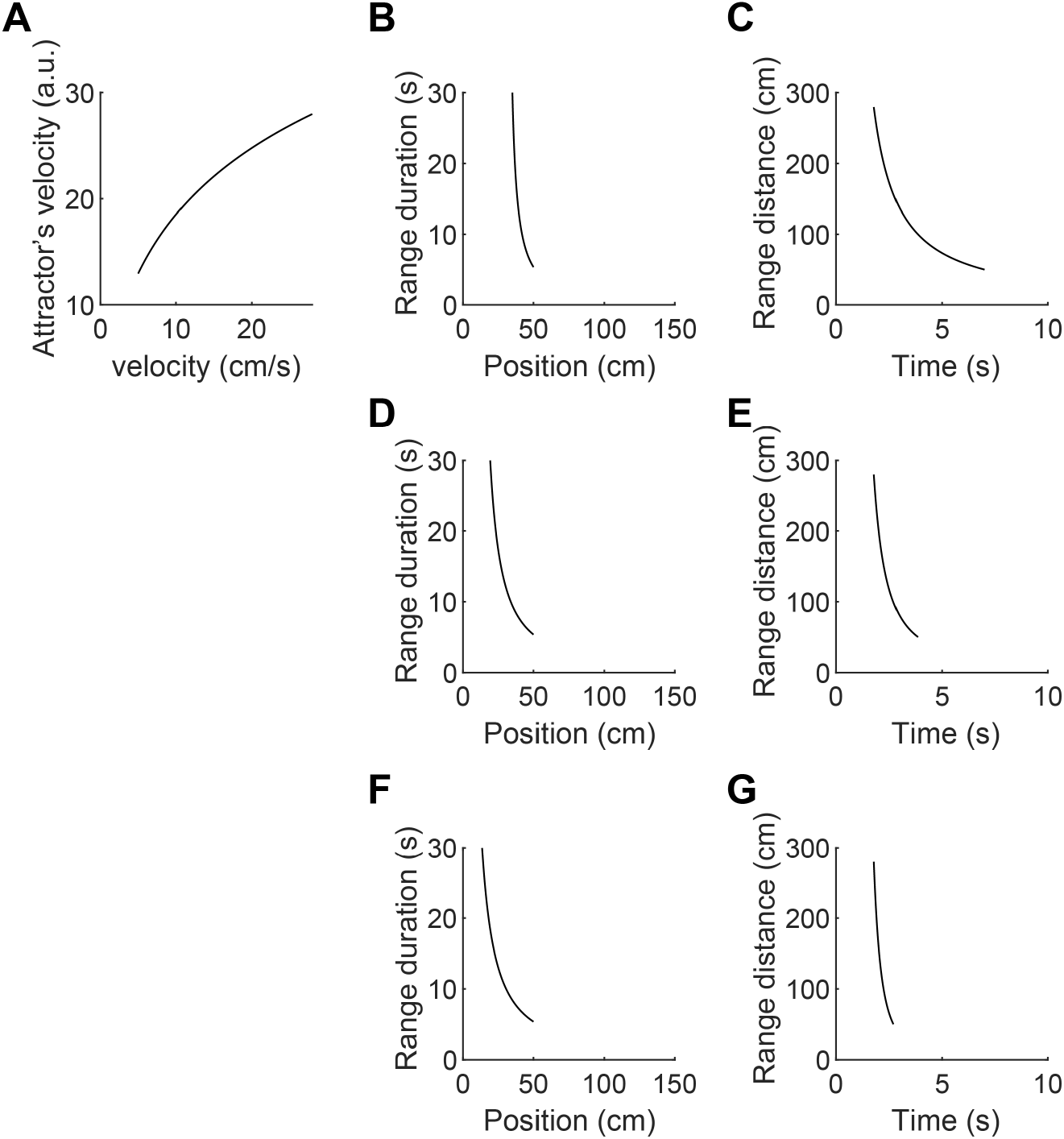
A continuous line attractor model that explains shifts in place and time fields. (**A**) Attractor’s velocity plotted against animal velocity (*v*), for 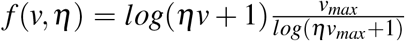, with *η* = 0.5 and *v*_*max*_ = 28. (**B-C**) Range duration as a function of the agent’s position (B), and range distance as a function of the elapsed time from the start of the lap (C) for *η* = 0.05 (neuron *n* = 50). (**D-E**) Idem (B-C) but for *η* = 0.5. (**F-G**) Idem (B-C) but for *η* = 5. As *η* increases, neurons tend to become more like time cells, and less like place cells.

**Figure 2.**
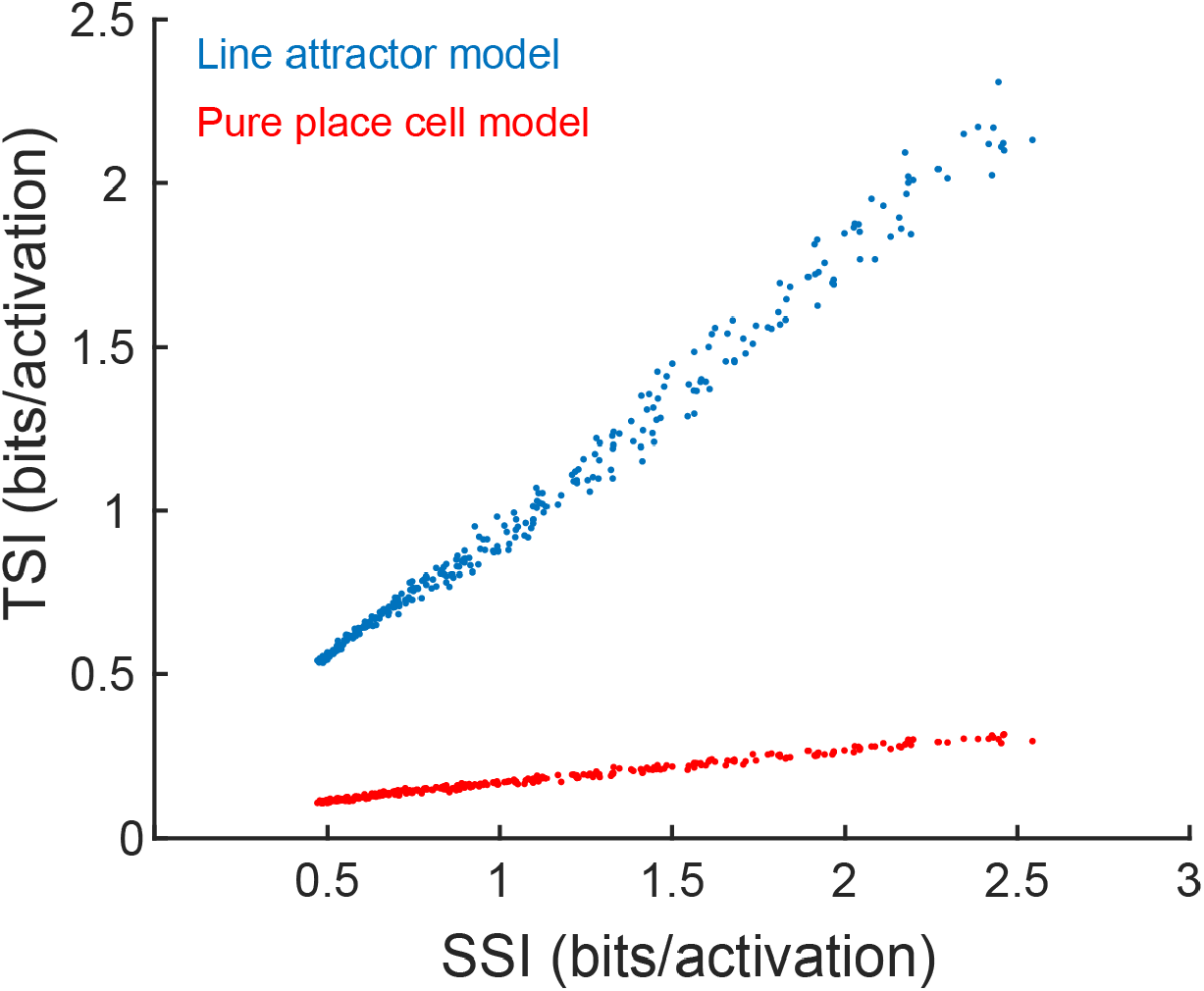
Temporal selectivity and spatial selectivity explained with a continuous line attractor with noise. Temporal selectivity is higher than the pure place cell model (place field combined with velocity). Neurons with higher variances have lower index values. A range of variances translates into a range of SSI and TSI values, which are positively correlated. The pure place field model has systematically lower TSI values because of the trade-off exemplified in Figure S1. *N* = 300 neurons (one dot per neuron).

**Figure 3.**
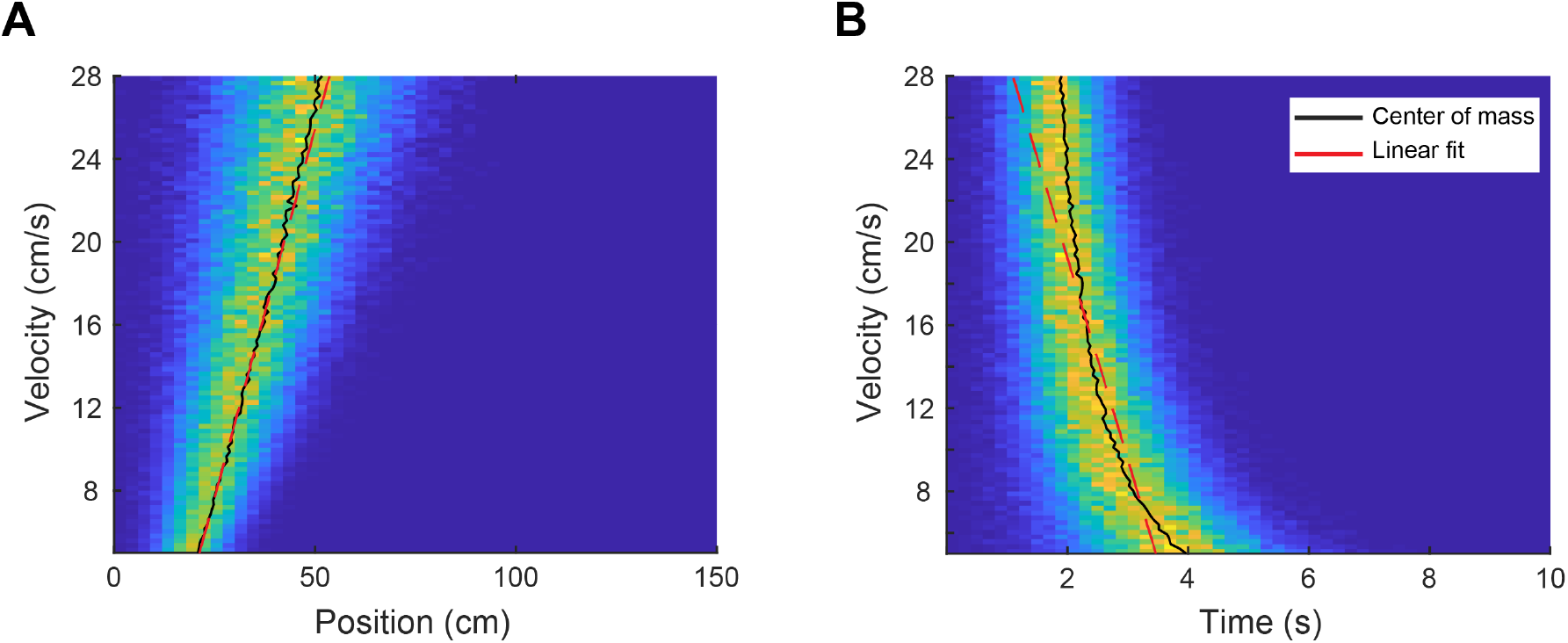
Non-linear shifts in the continuous line attractor. (**A**) Non-linear relationship of position with speed. A linear fit overestimate the centre of mass at maximum velocity. (**B**) Non-linear relationship of time with speed. A linear fit underestimates the centre of mass at maximum velocity.

**Figure 4.**
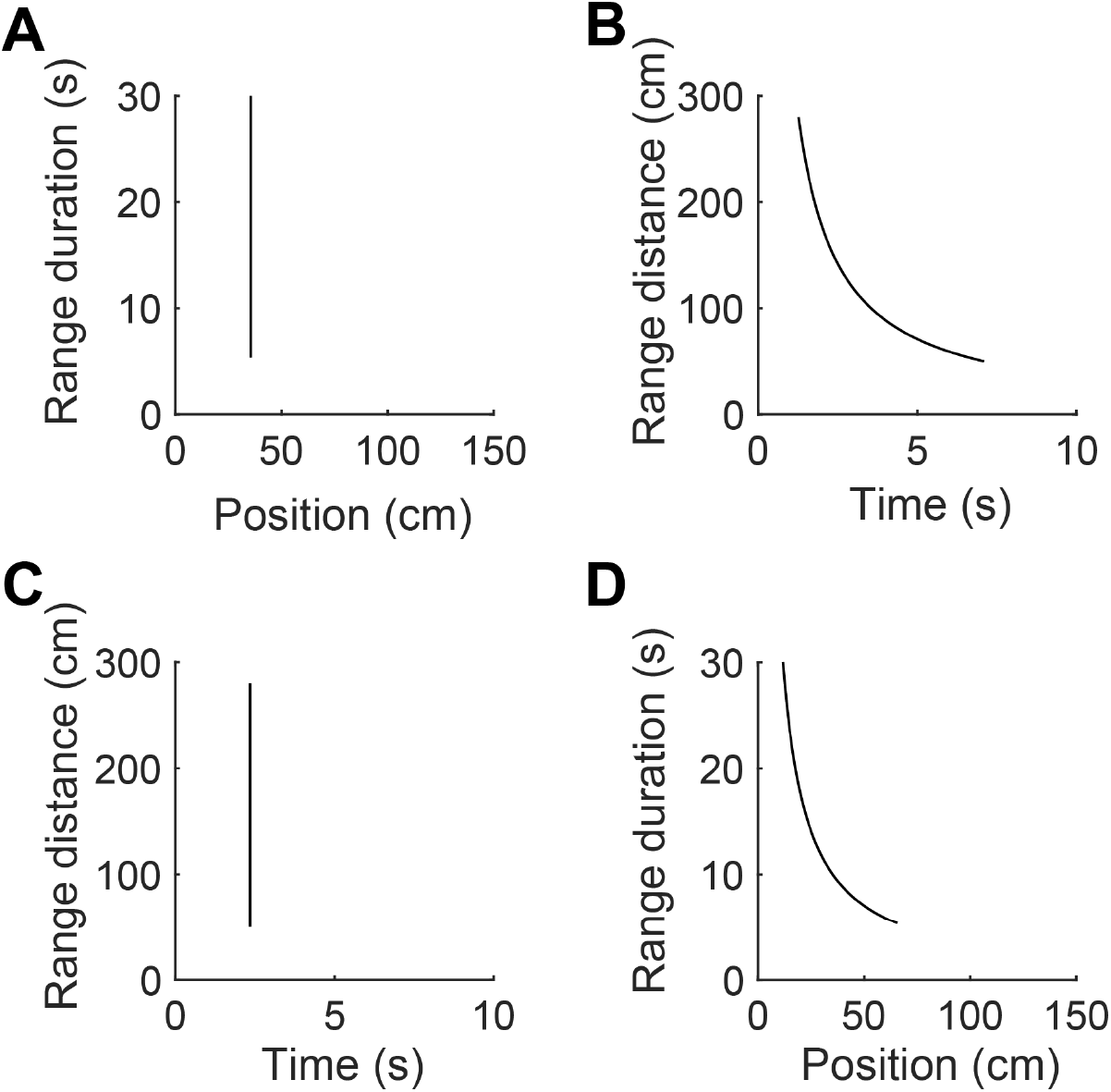
Space-Time trade-off in the Pure Place Cell Model. (**A**) Range duration for a neuron with a place field located at the mean encoded position by neuron in Fig. 1D (see Independence model in Chen et al., Figure 6). (**B**) The pure place cell model of panel A has less information about time than a neuron with mixed selectivity (compare with Fig. 1E). (**C**) Range duration for a neuron with a time field located at the mean encoded time by neuron in Fig. 1E. (**D**) The pure time cell model of panel C has less information about position than a neuron with mixed selectivity (compare with Fig. 1D).

In sum, our model reproduces the shifts in firing fields observed in Chen et al., the space-time trade-off, the observed differences in the space and time indices, the better fit of the space x time GLM, and the non-linear relationship of space and time with speed. The fact that a model that only integrates velocity explains key results from Chen et al. casts serious doubts regarding the conclusions of their study. In the line attractor model, the apparent time fields are in fact an artifact of the one-direction-only nature of the task. Analogously, the neurons found by Chen and colleagues could just be place cells, in which the coupling between animal speed and the attractor speed is a concave, non-decreasing, positive function. We are not suggesting that there are no time cells in Chen and colleagues’ study, nor that a space-time interaction is absent in the CA1 neural representations. We do believe, however, that the reported field shifts and space-time trade-off cannot be interpreted as clear signatures of the neural representation of a space-time continuum: more nuanced analyses are needed. First, a behavioural task that ensures complete decorrelation of time and space would be highly desirable, and could be attained by allowing animals to move in any direction, i.e.: back and forth in a linear track, or any direction in a 2D arena. Second, neural activity should be tested to falsify the null hypothesis instantiated by our model. This could be done at the level of individual neurons, and also at the neural population level, by embedding the population activity in a 1D manifold and assessing the rate of change within the manifold as a function of the animal’s speed. This function should be concave according to our model.

Concavity can be interpreted as the diminished capacity of response of the neural population to increasing speeds, stemming perhaps from the neuron’s limited dynamic range. It has already been found experimentally that speed cells’ firing rate is a concave function of animal’s speed[6]. Finally, other results in Chen et al., like the effect of CA3 inactivation and the shifts in multiple firing fields, fall outside the scope of our model, and may require a line attractor model with more neurophysiological detail.

## Methods

### Shifts in place and time fields explained with a continuous line attractor model

We will consider a mouse running in a linear track. The mouse has position *p* measured from the starting point, located at the origin of the track (*p*(*t* = 0) = 0). The position at time *t* is the solution to the integral 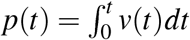, with *v*(*t*) > 0 the velocity of the agent. We will assume that, during a single run, the mouse’s velocity is constant, hence we have that *p*(*t*) = *vt*.

The neural representation of the mouse’s position can be modelled in terms of a continuous line attractor. This kind of attractor can be approximated by a neural network with a bump attractor dynamics. For simplicity, here we will consider a continuous linear attractor defined by 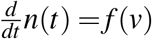. The variable *n*(*t*) encodes the active neuron at time *t*. Its derivative is a function of the animal’s velocity, linking changes in position with changes in the neural representation of this position. We will further assume that *n*(*t* = 0) = 0, and that *f* (*v*) is concave, positive and non-decreasing. Since *v* is constant, 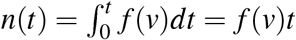. Therefore, a given neuron *n* will be activated at time

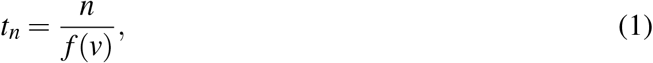

and the animal’s position during activation will be

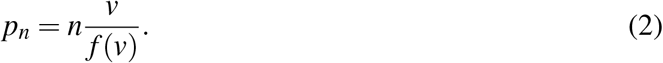

It is easy to see how velocity affects the timing of activation. As *v* increases, *t*_*n*_ decreases, replicating the shift towards earlier times seen in Chen et al., where laps are ordered in increasing velocity. To see the effect of velocity on the animal’s position at which *n* is activated, we derive *p*_*n*_ with respect to 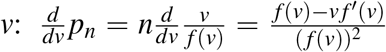, which is positive if *f* (*v*) − *v f* ^*′*^(*v*) > 0. By the mean value theorem, there exist *c ∈* (0, *v*) such that 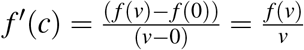. Since *f* ^*′′*^(*v*) < 0 (because *f* (*v*) is concave), then 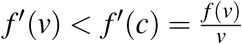 . Therefore, *v f* ^*′*^(*v*) < *f* (*v*), proving that *p*_*n*_ increases with *v*, replicating Chen’s results where place field shifted towards the end of the track when the animal run faster (decreasing range duration). If we further assume that *f* (*v*) is bounded, then *t*_*n*_ tends to a linear function of *n* as velocity increases, hence neurons tend to be perfect time cells in this scenario.

### Competition between space and time encoding

In the setting of Chen et al. the beginning of the lap is the origin from which both, position and time, are measured. Within the track, animals can only move forward from this frame of reference. This makes space and time utterly correlated. If we say that a neuron is both a place cell and a time cell, it means that it activates in a particular place and time from the lap start. This can only happen at a specific velocity (assuming a constant velocity throughout the lap). Conversely, if a neuron activates at a range of velocities (as in Chen et al. work), a trade-off between being pure place cell and pure time cell must be taking place.

To see this trade-off in the context of a line attractor, let’s consider function *f* described before (positive, concave and non-decreasing) and expand it by adding a parameter *η* which controls how much the function departs from the identity function. Let’s suppose it is also bounded. Specifically, we will assume that *f* (*η, v*) → *v* if *η* → 0 and *f* (*η, v*) → *c* if *η* → +∞, with *c* a positive constant. This in turn implies that 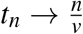 and *p*_*n*_ → *n* when *η* → 0, and that 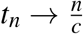 and 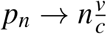 when *η* → +∞. In words, when the attractor’s velocity follows the animal’s velocity closer (low *η*), the position of activation becomes more independent of velocity, and the time of activation tends to be a linear function of the reciprocal of velocity (the neuron becomes more like a place cell). Conversely, as the coupling between the animal’s velocity and the attractor’s velocity becomes more sublinear (high *η*), the time of activation becomes more independent of velocity, and the position of activation tends to be a linear function of velocity (the neuron becomes more like a time cell).

In sum, the trade-off between space and time encoding follows from the experimental set-up in which space and time are necessarily correlated. Any simulated data analysis, like the one shown in Figure 6 of Chen et al. in which a neuron in considered to become a perfect place cell, or a perfect time cell, will necessarily lead to a drop in encoded information about the other dimension.

### Continuous line attractor implementation

For plots in Figure S1 we computed *p*_*n*_ and *t*_*n*_ according to eqs.1-2, taking

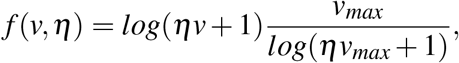

and setting *n* and a range of values for *η* to obtain curves that resemble some of the examples shown in Chen et al. The factor multiplying the logarithm is a constant that normalizes *f* output such that *f* (*v*_*max*_, *η*) = *v*_*max*_. This choice facilitates analysis, but has no critical role in the results obtained (the factor could be absorbed by the value of *n* in eqs. 1-2).

To facilitate comparison with Chen et al. figures, we plotted *p*_*n*_ (*x* axis “Position”) against “Range duration” (defined in Chen et al. as the time taken by the agent to cover 150 cm at speed *v*), and plotted *t*_*n*_ (*x* axis “Time”) against “Range distance (defined in Chen et al. as the distance covered by the animal after 10 s of moving at speed *v*).

### Line attractor with noise

We implemented a stochastic version of the above line attractor, in which a neuron activates with probability

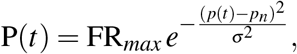

where *p*(*t*) is the position occupied by the animal at time *t, p*_*n*_ the position at which neuron *n* activates, according to the line attractor state, FR_*max*_ its maximum firing rate (expressed as activations per millisecond), and *σ* the parameter that regulates the width of the firing field. We simulated trajectories of 10 s (in 1 ms time steps), with velocity evenly distributed in the interval [*v*_*min*_ = 5 s, *v*_*max*_ = 28 s] (100 velocity values within the interval). We set *η* = 0.5, *σ* evenly distributed in the interval [20, 60] (30 values within the interval, 10 repetitions for each *σ* value), FR_*max*_ = 0.05 activations/ms and neuron *n* = 50. Average activations were computed in non-overlapping 100 ms windows. To compute the SSI and TSI values, we segmented space and time in 50 bins and calculated the mean activation within each bin. Then, the indices were defined as in Chen et al.: 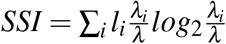, where *λ*_*i*_ is the mean activation of the ith bin, and *λ* the mean activation across bins, while *l*_*i*_ is the normalized occupancy of the ith spatial bin (in our case, all positions are visited equally). The TSI index was defined analogously, but time was binned in this case. We consider the activity of one neuron to be the pooled activity obtained across all velocities, for one repetition of sigma value. Therefore, we had 30 × 10 = 300 neurons. One SSI and one TSI were computed for each neuron (Figure S2).

We considered the model employed in Chen et al., that combines place fields with velocity (which we call “pure place cell model”, see Chen et al. Methods: “Predicted temporal selectivity from observed spatial selectivity”). This model instantiates the null hypothesis that an observed time field is the result of traversing across a place field at different velocities. For each simulated neuron in the line attractor we simulated a pure place cell neurons, with activation probability

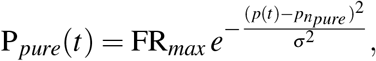

with 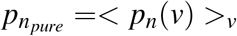 and <>_*v*_ symbolizing the average operation computed across velocities (see Figure 6. in Chen et al.).

The same data used to estimate the SSI and TSI were also employed to fit the neural activations with two separate Generalized Linear Models (GLMs), one that takes space and time as predictor variables, and another that takes space and velocity. Following Chen et al., we normalized position, time, and velocity to range between 0 and 1, and the peak activation rate was normalized to 15 Hz. The GLMs were fitted with the MATLAB function *fitglm*, with a fifth-order polynomial model and the Poisson distribution for the response variable.

The pure place cell model was also employed to compute the non-linear relationship of position and time with velocity (Figure S3). The velocity within the lowest 2/3 of the velocity range was used to fit a linear model to predict the centre of mass for the mean activation over time or position. Then, the linear model was employed to extrapolate the centre of mass at the highest velocity, which was compared to the actual centre of mass at the highest velocity.

For plots in Figure S4 we employed the pure place cell model, without noise, for the mean position and time encoded in Figure S1 D-E, to more clearly depict the space-time trade-off shown by this analysis.

## Data availability

The MATLAB code for generating all figures can be found at https://github.com/cmininni/simple_line_attractor.git

## Acknowledgements

This research was supported by the Consejo Nacional de Investigaciones Científicas y Técnicas Grant No. 11220200102316CO, Agencia Nacional de Promoción de la Investigación, el Desarrollo Tecnológico y la Innovación Grant No. PICT-2021-I-A-00957 and Universidad de Buenos Aires Grant No. 20020220200075BA.

## References

1. O’Keefe, J. Place units in the hippocampus of the freely moving rat. Experimental Neurology 51, 78–109. ISSN: 10902430 (1976).

2. MacDonald, C. J., Lepage, K. Q., Eden, U. T. & Eichenbaum, H. Hippocampal “time cells” bridge the gap in memory for discontiguous events. Neuron 71, 737–749. ISSN: 08966273. 10.1016/j.neuron.2011.07.012 (2011).

3. Howard, M. W. & Eichenbaum, H. Time and space in the hippocampus. Brain research 1621, 345–354 (2015).

4. Chen, S. et al. Integration and competition between space and time in the hippocampus. Neuron 112, 3651–3664. ISSN: 0896-6273. 10.1016/j.neuron.2024.08.007 (2024).

5. Khona, M. & Fiete, I. R. Attractor and integrator networks in the brain. Nature Reviews Neuroscience 23, 744–766. ISSN: 14710048. arXiv: 2112.03978 (2022).

6. Góis, Z. H. T. D. & Tort, A. B. L. Characterizing speed cells in the rat hippocampus. Cell reports 25, 1872–1884 (2018).

